# xTracer: Integrating chromatogram and mobiligram correlations for untargeted peptide identification in SLIM-based PAMAF data

**DOI:** 10.64898/2026.02.11.705440

**Authors:** Jian Song, Liulin Deng, Lauren Royer, Bennett Kalafut, Daniel DeBord, Jesse G. Meyer

## Abstract

Parallel Accumulation with Mobility-Aligned Fragmentation (PAMAF) achieves near-complete ion utilization and high spectral specificity by fragmenting all mobility-separated precursors without quadrupole isolation. Leveraging the ultrahigh mobility resolution of SLIM, this quadrupole-free strategy maximizes ion utilization efficiency and offers a promising approach in mass spectrometry–based proteomics, particularly for low-abundance peptides or low-input samples. However, the unique data structure of PAMAF where precursor–fragment relationships are encoded along the mobility dimension renders it incompatible with existing peptide identification tools. Here, we present xTracer, the first untargeted peptide identification algorithm developed specifically for PAMAF data. xTracer integrates correlations across both chromatographic and mobility dimensions to associate precursor and fragment ions, reconstruct pseudo-spectra, and enable database searching using well-established DDA search engines. Applied to datasets with varying sample loads and acquisition throughputs, xTracer consistently achieved robust and reproducible peptide identifications, outperforming single-domain correlation strategies. Overall, xTracer provides a versatile and high-efficiency computational framework for reconstructing pseudo-spectra from quadrupole-free, mobility-aligned fragmentation data, enhancing the analytical power of high-resolution ion mobility (HRIM)–based proteomics.

## 1 Introduction

Tandem mass spectrometry (MS/MS) has become the analytical foundation of proteomics, offering the molecular specificity required for confident peptide and protein identification^[1]^. In conventional implementations of MS acquisition, peptide ions are isolated by a quadrupole filter prior to fragmentation and detection, either through narrow isolation windows in data-dependent acquisition^[2]^ (DDA) or wider windows in data-independent acquisition^[3]^ (DIA). While quadrupole-based isolation provides high precursor selectivity, it is inherently inefficient, as ions outside the isolation window are discarded, severely limiting ion utilization efficiency^[4]^. For instance, the estimated ion utilization efficiency of typical DDA or narrow-window DIA schemes using a 2 Th isolation window is approximately 1%^[4, 5]^, whereas a conventional 32-window DIA design achieves only about 3%^[5, 6]^. The diaPASEF method, which synchronizes quadrupole isolation windows with trapped ion mobility separation, improves ion utilization by approximately fivefold compared to conventional DIA while maintaining spectral specificity^[6]^. Nevertheless, the absolute ion utilization remains low—around 15%—indicating that even mobility-assisted quadrupole isolation still discards the majority of ions. Inevitably, current isolation strategies that rely on a quadrupole suffer from low ion utilization efficiency, which poses a major obstacle for the identification of low-abundance peptides and low-input samples.

One promising approach to overcoming these limitations is Parallel Accumulation with Mobility Aligned Fragmentation (PAMAF), a quadrupole-free fragmentation strategy based on high-resolution ion mobility separations^[7, 8, 9]^. PAMAF separates precursors in the ion mobility dimension rather than the *m/z* dimension using structures for lossless ion manipulation^[10, 11, 12]^ (SLIM), which currently provides the highest ion mobility resolution available. Supported by high-speed mass spectrometry acquisition (approximately 150 µs per pulse for time-of-flight acquisition), nearly all precursors separated by SLIM can be captured and fragmented, achieving close to 100% ion utilization without discarding ions. The resulting fragment ions are temporally aligned with their corresponding precursors in mobility space, enabling comprehensive fragmentation through continuous scanning in the ion mobility dimension rather than sequential quadrupole scanning (Fig. 1a). This approach dramatically improves ion utilization efficiency and, through the high resolution of SLIM, prevents spectra from becoming overly congested or chimeric due to coeluting isobars. However, the data structure of PAMAF fundamentally differs from traditional DIA data: precursor–fragment relationships are encoded along the mobility axis, and no explicit *m/z* isolation boundaries exist. Consequently, the novel data structure of PAMAF limits the applicability of existing peptide identification software, and its analytical potential in proteomics remains largely unexplored.

**Fig. 1.**
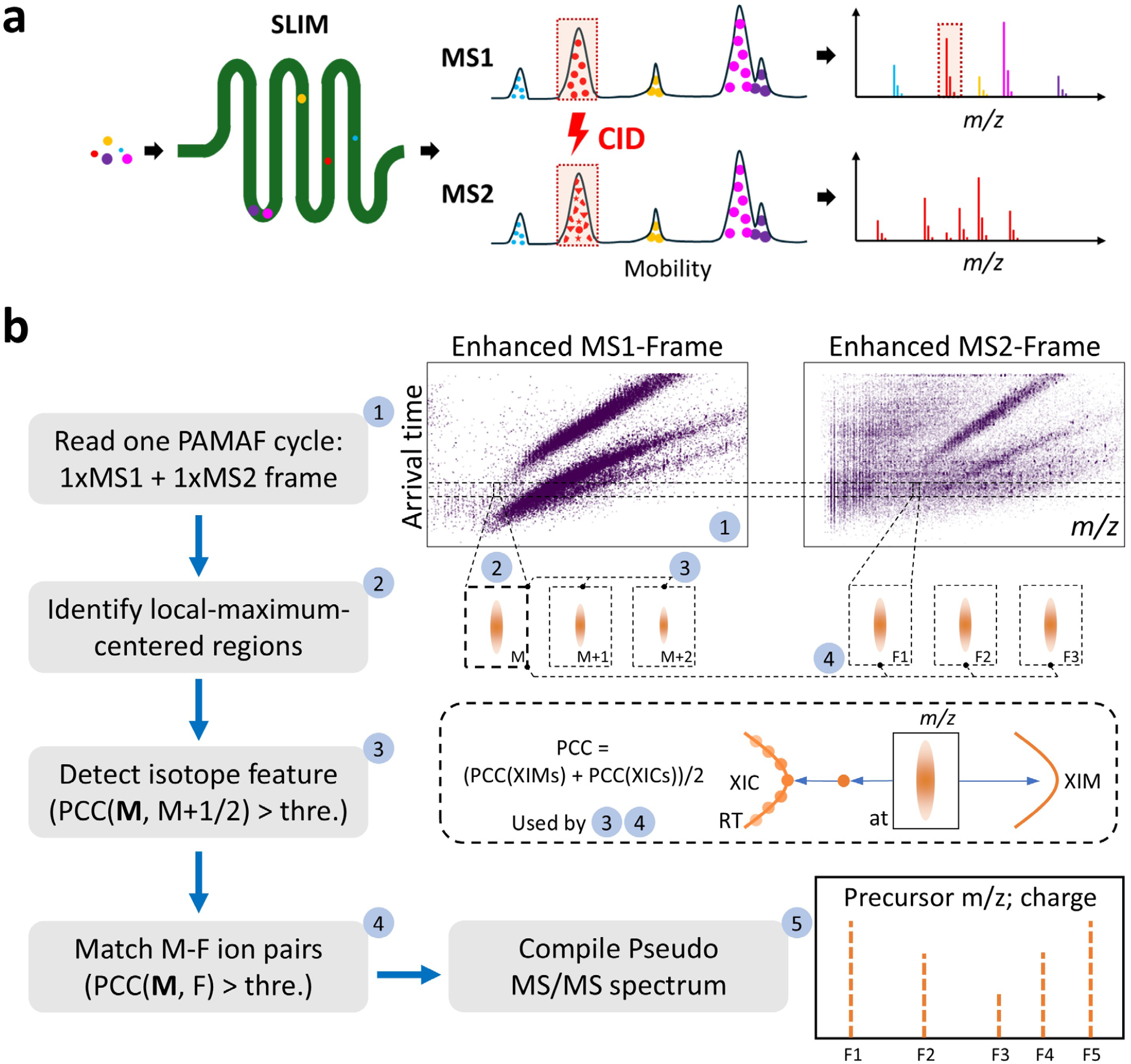
Overview of the PAMAF principle and the xTracer workflow. **a)** Schematic illustration of the PAMAF (Parallel Accumulation with Mobility-Aligned Fragmentation) principle, highlighting quadrupole-free fragmentation, near-complete ion utilization, and mobility-aligned ion dissociation. **b)** Overview of the xTracer workflow for untargeted peptide identification, including local maxima detection, extraction of chromatograms (XICs) and mobiligrams (XIMs), isotope feature detection, and correlation-based precursor–fragment pairing.

In the context of DIA data analysis, peptide identification strategies can be broadly categorized as targeted or untargeted, which often correspond to peptide-centric and spectrum-centric methodologies, respectively^[13]^. Targeted approaches depend on pre-existing spectral libraries to extract peptide signals, limiting their applicability when such libraries are unavailable or incomplete^[14, 15]^. In contrast, untargeted strategies operate without prior spectral information, reformulating DIA data into collections of pseudo-spectra that resemble those obtained from DDA^[16]^. These pseudo-spectra can then be searched using well-established DDA search engines, rendering untargeted identification particularly advantageous for novel data types such as PAMAF. DIA-Umpire^[16]^ was the first untargeted algorithm developed for DIA data. It infers precursor– fragment relationships by correlating chromatograms, thereby grouping co-eluting fragment ions with their corresponding precursors to construct pseudo-spectra. MSFragger-DIA^[17]^, in contrast, first filters potential peptide candidates for each chimeric DIA spectrum and subsequently assigns fragment peaks to the remaining candidates to achieve deconvolution. diaTracer^[18]^ is, to date, the only software capable of performing spectrum-centric identification for ion mobility–enhanced DIA data. It identifies local maxima within the ion mobility-*m/z* plane, performs the centroiding, extracts the corresponding chromatograms, and evaluates their correlations to pair precursor and fragment ions, thereby generating pseudo-spectra in a manner analogous to DIA-Umpire.

Inspired by diaTracer, we developed xTracer, an untargeted peptide identification algorithm tailored for SLIM-based PAMAF data. Given the ultrahigh ion mobility resolution of SLIM, xTracer incorporates correlations from both chromatograms and mobiligrams to optimize precursor– fragment pairing, thereby improving the accuracy of pseudo-spectra generated directly from PAMAF data. Applied across datasets with varying sample loads and LC gradients, xTracer consistently achieved robust identification, yielding thousands of peptide sequences from standard HeLa samples. Comparative analyses further demonstrated that integrating chromatogram- and mobiligram-based correlations is helpful for precursor–fragment association compared with using either domain alone. This work represents the first successful demonstration of untargeted peptide identification for PAMAF data. The resulting peptide identifications can be used to construct spectral libraries, thereby supporting the development of future targeted analyses. Overall, xTracer provides a general computational framework for interpreting quadrupole-free, high-resolution mobility-enabled fragmentation spectra, establishing PAMAF as a viable operating mode for comprehensive proteome profiling.

## 2 Materials and Methods

### Experimental Conditions

#### Sample preparation

In this study, a Pierce HeLa Protein Digest Standard (P/N, 88328, Thermo Fisher Scientific, Waltham, MA) was used. The material was received as a lyophilized powder, reconstituted in water containing 0.1% formic acid (v/v), and serially diluted to the desired concentration depending on the target sample load. To enable potential retention time calibration for the HeLa samples analyzed by liquid chromatography (LC), indexed Retention Time (iRT) Calibration Peptides (Biognosys iRT Kit, P/N 1816351, Bruker, Billerica, MA) were spiked into all samples. Various sample loads (25, 50, and 100 ng HeLa on column) were analyzed in triplicate across the various chromatographic methods.

#### Instrumentation

All LC analyses were performed on an Evosep One UHPLC system (Evosep, Odense, Denmark). Samples were prepared on Evotips using the standard Evosep protocol, and all separations employed the preconfigured Evosep methods without modification. Mobile phase A was water with 0.1% formic acid (v/v), and mobile phase B was acetonitrile with 0.1% formic acid (v/v). Data were acquired at various sample throughput settings while maintaining the same system operating mode. For the 60 and 100 samples-per-day (SPD) methods, peptides were separated on an 8 cm x 150 µm i.d. reversed-phase column packed with 1.5 µm C18-coated porous silica beads heated to 40°C (Evosep EV 1109). For the 200 SPD method, peptides were separated on a 4 cm x 150 µm i.d. reversed-phase column packed with 1.9 µm C18-coated porous silica beads (Evosep EV1107). The outlet of each LC column was connected to an Evosep EV1116 Agilent Sleeve Adapter fitted with a 30 µm stainless steel nanospray emitter (Evosep EV1086), mounted in a custom holder and positioned in front of the capillary inlet of the high-resolution ion mobility/quadrupole-time-of-flight mass spectrometer (HRIM/Q-TOF MS).

A prototype (HRIM/Q-TOF) system was used for all analyses. The HRIM device is based on a MOBIE system (MOBILion Systems, Chadds Ford, PA) coupled with an Agilent 6546 Q-TOF mass spectrometer (Agilent Technologies, Santa Clara, CA). The instrument has been described in detail previously^[7, 8, 9]^. Briefly, ions are separated by HRIM in the SLIM module and then transmitted to the Q-TOF MS for either precursor mass analysis or collision-induced dissociation (CID) followed by fragment mass analysis. In the parallel accumulation mobility-aligned fragmentation (PAMAF) operating mode, ion mobility frames alternate between low and high collision energies (CE); fragment ions detected in a given high energy frame correspond to precursor ions accumulated and separated in the preceding low energy frame.

### xTracer workflow

The workflow of xTracer is illustrated in Fig. 1b. After loading the PAMAF file using the MOBILion software development kit (MBI-SDK) for reading the raw .mbi file format, the algorithm sequentially processes each cycle in the PAMAF data. Each cycle contains one MS1 frame and one MS2 frame, and each frame can be regarded as a collection of ions characterized by their arrival time, *m/z*, and intensity, where the arrival time reflects the ion mobility separation. xTracer first sums the signals of the current cycle with those of its neighboring cycles to enhance the signal intensity, generating enhanced MS1 and enhanced MS2 frames. It then detects local maxima in the enhanced frames. A local maximum is defined as a point that (i) has the highest intensity within its surrounding region (defined by ± *tol_at_area* ms in arrival time and ± *tol_ppm* ppm in *m/z*), and (ii) has at least a minimum number of supporting signal points (≥ *tol_point_num*). For each detected local maximum, xTracer extracts both its chromatogram and mobiligram. Specifically, the chromatogram spans multiple cycles (default = 7 cycles centered on the current one), where each data point represents the summed intensity within the tolerance region. The mobiligram, in contrast, is derived from the current enhanced cycle by projecting all signals within the tolerance region onto the mobility axis, forming a vector representation (default: 15 bins). xTracer next performs isotopic feature detection on all local maxima in the MS1 frame. If a given local maximum has two additional local maxima whose *m/z* values follow the isotopic pattern (corresponding to M, M+1, and M+2), with arrival-time deviations within *tol_at_shift* ms, and if their correlation (defined as the average of the chromatogram and mobiligram Pearson correlation coefficients, PCCs) exceeds a defined threshold (*tol_pcc*), the peak is regarded as a potential precursor. After determining the potential precursors, xTracer searches the MS2 frame for local maxima whose arrival time deviation from the precursor is within *tol_at_shift* ms. If the correlation between the two signals (averaged over chromatogram and mobiligram PCC correlations) exceeds *tol_pcc*, the MS2 ion is considered a potential fragment ion of that precursor. When more than *tol_fg_num* fragment ions are associated with a precursor, xTracer reconstructs a pseudo-spectrum containing the retention time, arrival time, fragment *m/z*, and the summed intensity within the tolerance window. Finally, all reconstructed pseudo-spectra are exported in the .mgf text format.

The .mgf files generated by xTracer were subsequently searched using the DDA search engine Sage^[19]^ (version 0.15.0-beta.1) for peptide identification. Search parameters were kept as close to the default settings as possible (see Supplement Fig. 1). The searches were performed against a *Homo sapiens* FASTA database (taxon identifier: 9606; 20,420 reviewed sequences) downloaded from UniProt (June 2024 release). To determine the generally applicable parameters of xTracer, we compared peptide identification results from Sage under 60 random different xTracer parameter combinations on two datasets (see Supplementary Fig. 2). Based on the top-3 combinations for each dataset result, xTracer empirically uses the following as the default parameters: *tol_at_area* = 2 ms, *tol_at_shift* = 1 ms, *tol_ppm* = 30, *tol_pcc* = 0.4, *tol_point_num* = 5 and *tol_fg_num* = 10.

## Data and Code Availability

Raw PAMAF MS data, xTracer results, and Sage search outputs have been deposited in the MassIVE repository (https://massive.ucsd.edu; project MSV000099577). The xTracer software is publicly available at https://github.com/xomicsdatascience/xtracer and can be installed via “*pip install xtracer-pamaf*”. The github repository also provides information about accessing the MBI-SDK to read the PAMAF data.

## 3 Results and Discussion

Because xTracer computes ion correlations by jointly considering chromatograms and mobiligrams, we compared its performance with two derivative algorithms that use only a single correlation domain: xTracer-XIC (computing PCC using chromatograms only) and xTracer-XIM (computing PCC using mobiligrams only).

In the variable loading dataset (Fig. 2), all three methods generated pseudo-spectra far exceeding the number of spectra expected from DDA under equivalent conditions (assuming a 50 Hz acquisition rate and a 45 min gradient, DDA will yield ∼100,000 spectra). Among them, the number of pseudo-spectra ranged from approximately 2–12x that of DDA, with xTracer-XIM consistently producing the highest numbers, xTracer-XIC the fewest, and xTracer falling in between. Increasing sample load led to a higher number of pseudo-spectra and an increase in the number of fragment ions per pseudo-spectrum. At the fragment level, the PCC of fragment ions tended to increase with their intensity, indicating that higher-intensity fragments were more reliably captured in the pseudo-spectra. Comparing the three methods, xTracer-XIM appeared to generate pseudo-spectra in a relatively permissive manner, resulting in a larger number of spectra with more fragment ions per spectrum; xTracer-XIC, in contrast, produced the fewest pseudo-spectra and fragment ions; and xTracer displayed intermediate spectrum and fragment counts. Interestingly, the ranking based on fragment-level PCC was xTracer-XIC > xTracer-XIM > xTracer, which may be because xTracer, by jointly considering both chromatographic and ion mobility constraints, includes additional borderline fragments supported by only one dimension, slightly lowering the overall PCC despite maintaining a moderate number of fragment ions. After Sage searches, xTracer generally achieved the highest numbers of identified peptide spectrum matches (PSMs, at 1% PSM-level FDR) and peptides (at 1% peptide-level FDR), followed by xTracer-XIM and xTracer-XIC. At 25, 50, and 100 ng, xTracer showed −4.0%/6.0%/13.0% and 15.0%/11.0%/6.0% changes in peptide identifications relative to xTracer-XIM and xTracer-XIC, respectively (based on the total identifications across three replicates), with Wilcoxon rank-sum p-values of 0.80/0.20/0.05 and 0.35/0.05/0.05. The overall spectrum identification rate was approximately 10%, and identified peptides were largely overlapping among the three methods (Fig. 3a), with no significant difference in reproducibility across replicates. xTracer also demonstrated high consistency and reliable detections across different sample loads (Fig. 3c): 99.8% of peptides detected in the 25 ng sample were found in the 50 ng sample, 99.8% of peptides from the 50 ng sample were observed in the 100 ng sample, and 96.3% of peptides identified in the 100 ng sample were covered by the in-house fractionated DIA library (∼120,000 peptides). The fraction of matched b/y ions relative to theoretical ions was similar (∼30%) for all methods. While this is lower than that observed for DDA (∼40–50%) it is consistent with prior DIA-Umpire findings^[16]^. This reduction likely reflects the suppression of low-intensity fragment ions by coexisting high-intensity species within the same SLIM scan and the incomplete deconvolution of coeluting precursors. In terms of computational efficiency (tested on a laptop with a 13th Gen Intel Core i9-13950HX CPU and 64 GB of RAM), xTracer-XIM was the fastest, xTracer the slowest, and xTracer-XIC intermediate (Supplementary Fig. 3). Processing time increased with sample load but remained well below the MS acquisition time in all cases.

**Fig. 2.**
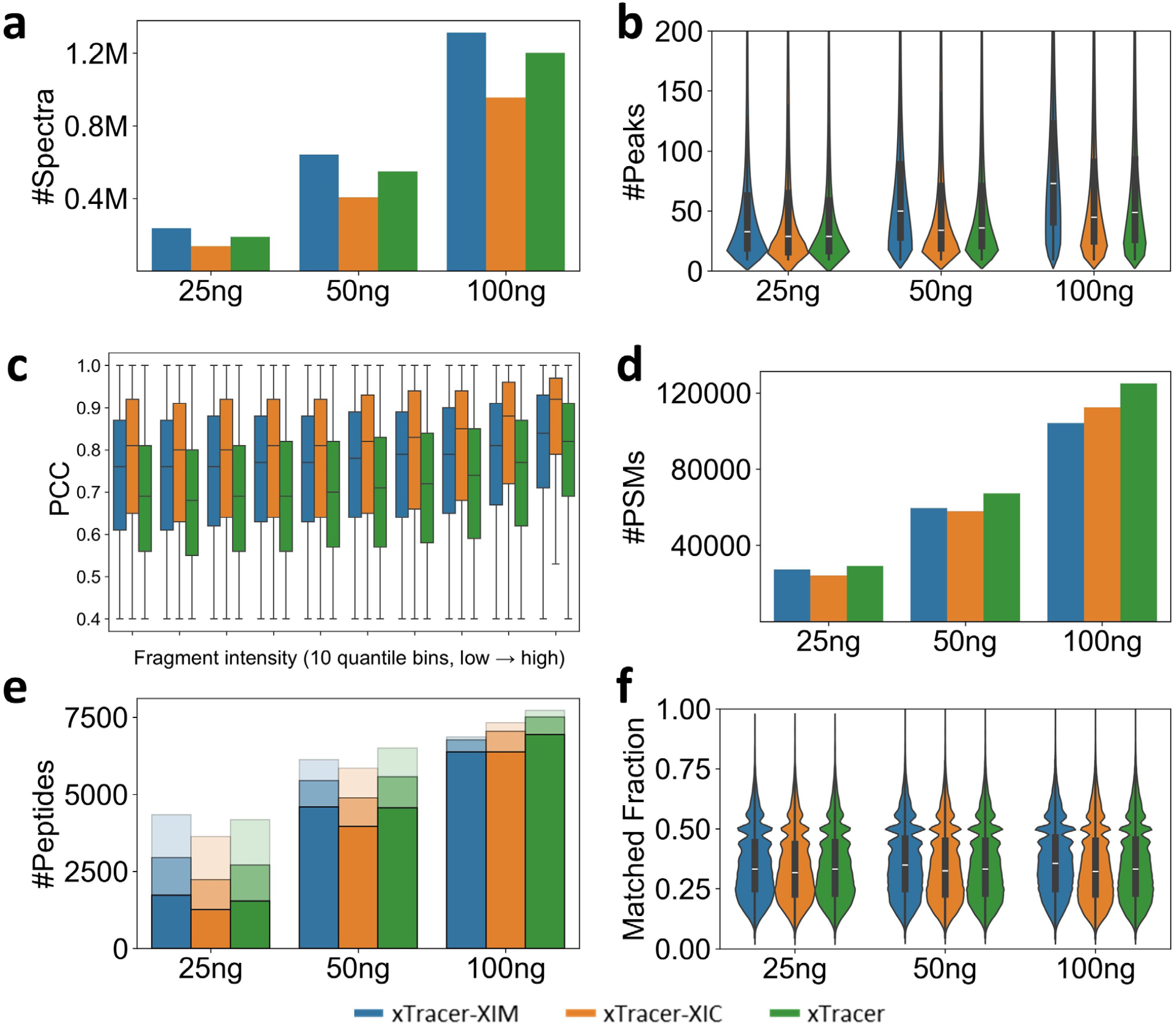
Performance of xTracer across different sample loads. xTracer-XIM denotes correlation computation using only mobiligrams, xTracer-XIC uses only chromatograms, and xTracer combines both by averaging correlations from the two domains. **a)** Average number of reconstructed pseudo-spectra from triplicate runs. **b)** Distribution of the number of fragment ions per pseudo-spectrum. **c)** Box plots showing the relationship between fragment-ion intensities and PCCs in the generated pseudo-spectra from 50 ng samples. Fragment ions are binned into 10 intensity quantiles from low to high. **d)** Average number of PSMs identified at 1% FDR. **e)** Average number of peptides identified at 1% FDR. The bar charts represent identification counts from triplicate injections, with colors from light to dark indicating detection in one, two, or all three replicates. **f)** Fraction of fragment ions matched within identified PSMs under 1% FDR.

**Fig. 3.**
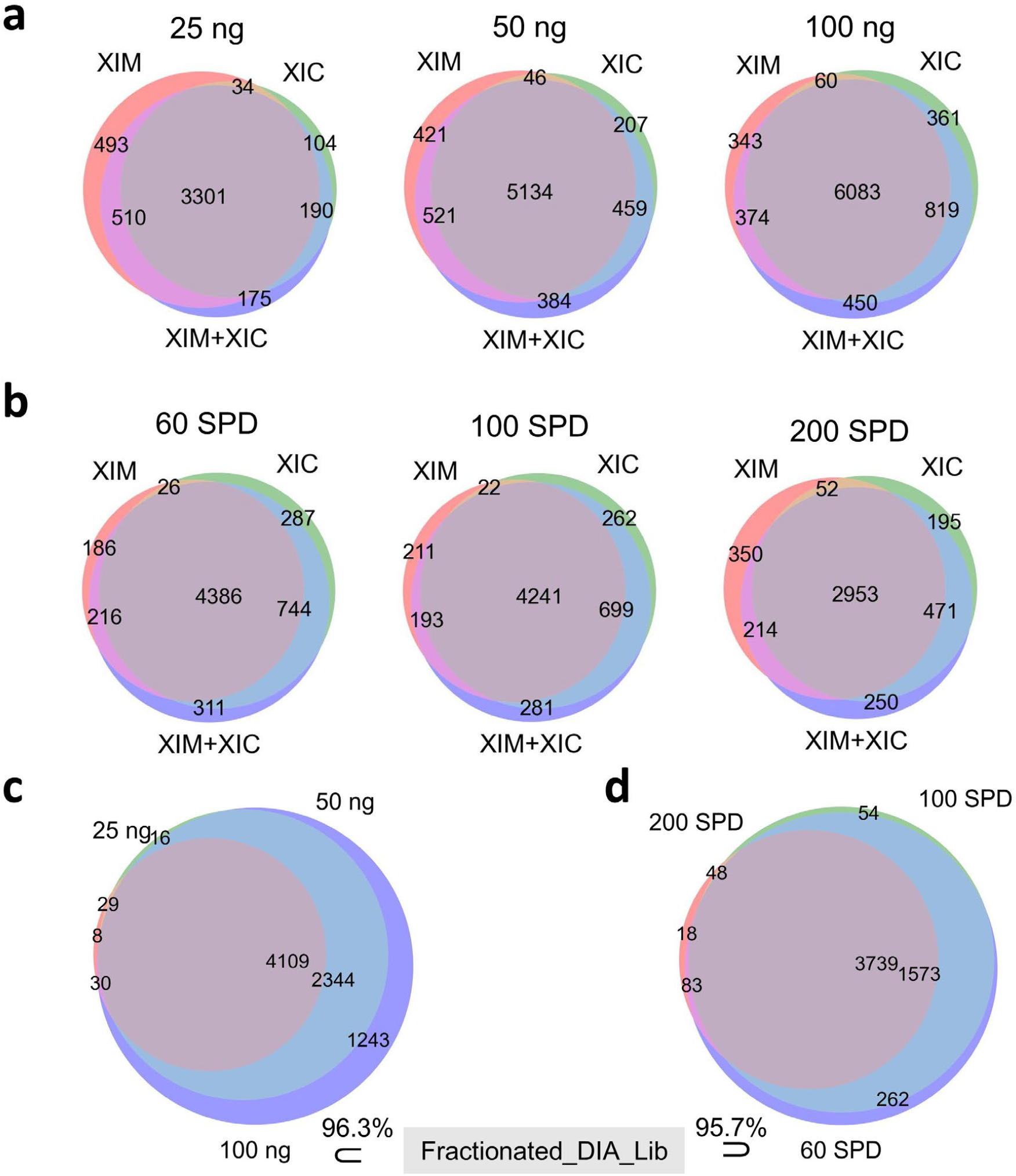
Overlap of peptide identifications. xTracer-XIM (labeled as “XIM” in the figure) uses only mobiligram correlations, xTracer-XIC (“XIC”) uses only chromatogram correlations, and xTracer (“XIM+XIC”) combines both domains. **a)** Venn diagrams showing peptides identified at 1% FDR from datasets with different sample loads. **b)** Venn diagrams showing peptides identified at 1% FDR from datasets acquired at different throughputs. **c)** Overlap of peptide identifications across different sample loads using xTracer. 96.3% of the peptides identified in the 100 ng sample are verified by the fractionated DIA library (approximately 120,000 peptides). **d)** Overlap of peptide identifications across different throughputs using xTracer. 95.7% of the peptides identified in the 60 SPD sample are verified by the fractionated DIA library.

In the variable throughput dataset (Fig. 4), similar patterns were observed. xTracer-XIM again produced the highest numbers of pseudo-spectra and peaks, xTracer-XIC the lowest, and xTracer intermediate. The fragment-level PCC of xTracer remained the lowest among the three methods. Similarly, higher throughput led to increased signal density, which in turn resulted in a corresponding rise in the total number of peaks. Following Sage identification, xTracer again yielded more PSMs and peptide identifications than xTracer-XIM and xTracer-XIC. At 60, 100, and 200 SPD, xTracer showed 18.0%/16.0%/9.0% and 4.0%/4.0%/6.0% changes in peptide identifications relative to xTracer-XIM and xTracer-XIC, respectively (based on the total identifications across three replicates), with Wilcoxon rank-sum p-values of 0.05/0.05/0.04 and 0.35/0.05/0.04. The three methods showed high overlap in peptide identifications (Fig. 3b) and comparable reproducibility across replicates. The spectrum identification rate (∼5–10%) and the ion matching ratio (∼30%) were consistent with those observed in the variable loading experiment. Consistently, peptide identifications by xTracer were largely covered across different throughput levels (Fig. 3d): 99.5% of peptides identified in the 200 SPD sample were also found in the 100 SPD sample, 99.0% of peptides from the 100 SPD sample were observed in the 60 SPD sample, and 95.7% of peptides from the 60 SPD sample were included in the fractionated DIA library. These results generally support the accuracy of the identifications. For analysis time, xTracer-XIM maintained the shortest processing time, followed by xTracer-XIC and xTracer, and overall processing became faster at higher throughput, yet remained substantially shorter than data acquisition.

**Fig. 4.**
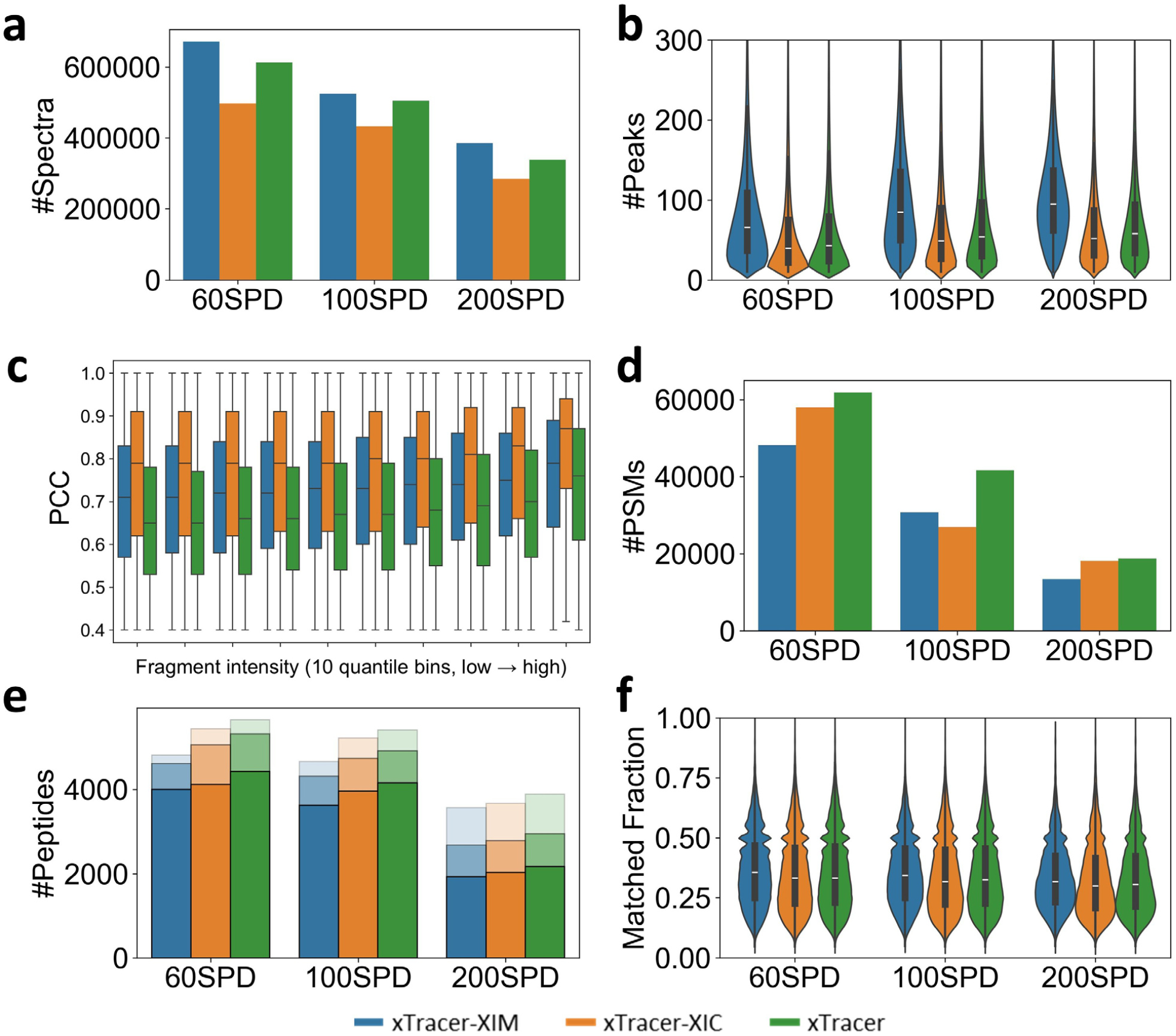
Performance of xTracer across datasets with varying acquisition throughputs. xTracer-XIM denotes correlation computation using only mobiligrams, xTracer-XIC uses only chromatograms, and xTracer combines both by averaging correlations from the two domains. **a)** Average number of reconstructed pseudo-spectra from triplicate runs. **b)** Distribution of the number of fragment ions per pseudo-spectrum. **c)** Box plots showing the relationship between fragment-ion intensities and PCCs in the generated pseudo-spectra from 100 SPD samples. Fragment ions are binned into 10 intensity quantiles from low to high. **d)** Average number of PSMs identified at 1% FDR. **e)** Average number of peptides identified at 1% FDR. The bar charts represent identification counts from triplicate injections, with colors from light to dark indicating detection in one, two, or all three replicates. **f)** Fraction of fragment ions matched within identified PSMs under 1% FDR.

Collectively, results from both datasets reveal several insights:

1. Pseudo-MS/MS spectra extraction along mobility and time is effective for untargeted peptide identification from PAMAF data. This is incredible despite the absence of precursor filtering and the fragmentation of all ions resulting from the omission of quadrupole selection.
2. xTracer-XIM consistently produced more pseudo-spectra and peaks than xTracer-XIC. This likely arises because mobiligram-based correlations are computed within single DIA cycles, whereas chromatogram-based correlations span multiple cycles. The noise from any single cycle can introduce deviations in the chromatogram-based correlations. Hence, PCC values derived from XIM more readily exceed identical thresholds.
3. The number of pseudo-spectra does not directly correlate with the number of identified peptides. Excessive pseudo-spectra generation can increase the overall false positive proportion, leading to more conservative FDR control and consequently fewer identifications^[20]^.
4. xTracer generally outperformed xTracer-XIC and xTracer-XIM in peptide identifications, underscoring the benefit of combining chromatogram- and mobiligram-based correlations. As illustrated in Fig. 5, for the identified precursor by xTracer, some fragment ions exhibited high PCCs by mobiligrams but low PCCs by chromatograms, while others showed the opposite. By integrating both correlations, more fragment ions can be correctly paired with their precursors, thereby increasing the confidence of peptide identification. This case clearly demonstrates the complementarity between the chromatogram and mobiligram domains under the high ion-mobility resolution conditions enabled by SLIM.

**Fig. 5.**
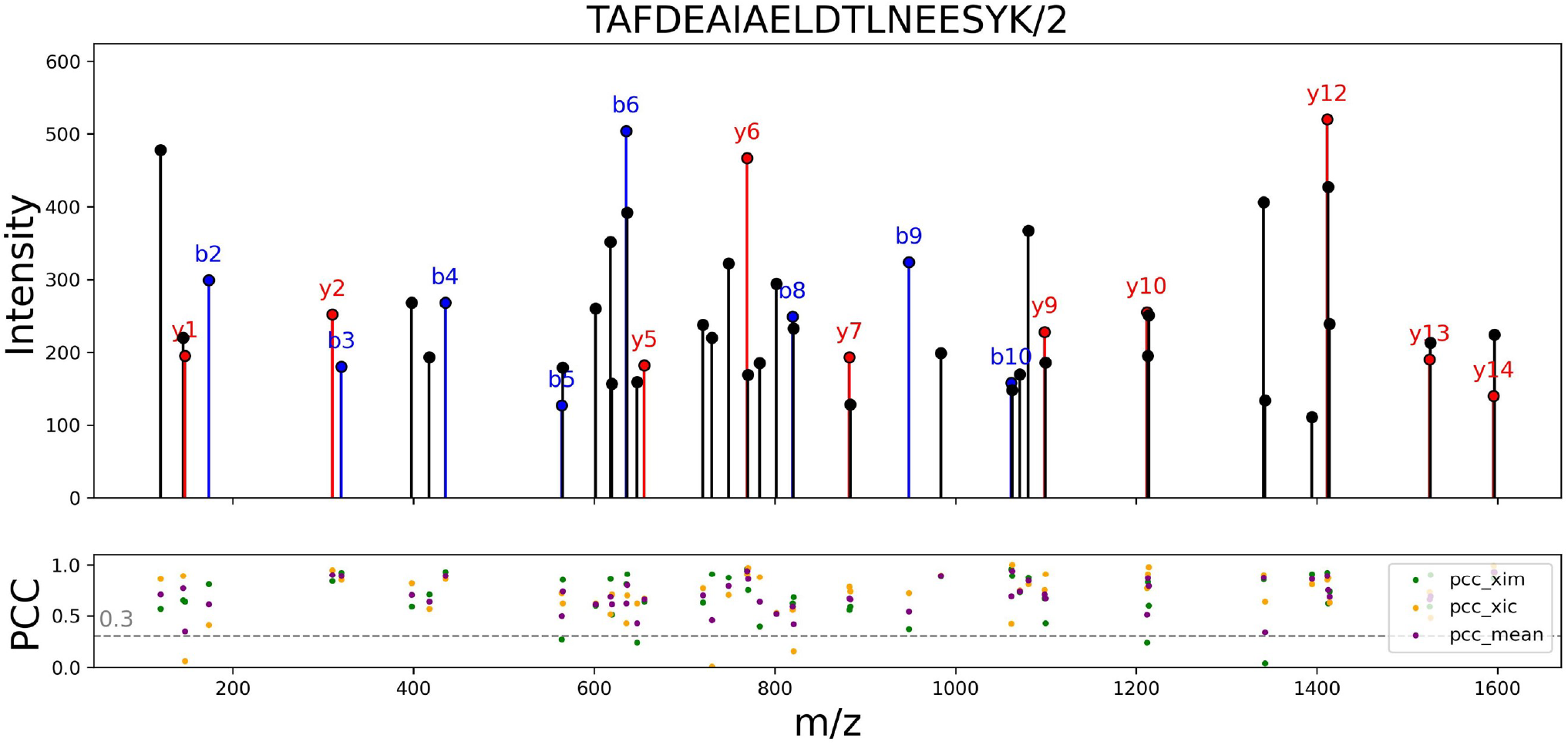
Case study illustrating the peptide pseudo-spectrum matches. The upper panel shows a representative pseudo-spectrum generated by xTracer and the identified precursor, along with the matched b/y fragment ions. The lower panel displays the fragment–precursor correlations computed based on chromatogram (pcc_xic), mobiligram (pcc_xim), and their average (pcc_mean).

The peptide identification results obtained by xTracer from the above datasets indicate that, although the algorithm jointly evaluates chromatogram and mobiligram correlations, it has not yet translated the near-complete ion utilization efficiency of PAMAF into deep proteome coverage. Similar to other untargeted analysis frameworks such as DIA-Umpire, pseudo-spectrum generation in xTracer primarily depends on the presence of well-defined precursor features in the MS1 domain, as peptide identification is initiated from detected MS1 isotope features. However, the relatively high coupling of ion signals in MS1 can introduce interference and potentially lead to peptide misidentifications. Future improvements should therefore focus on relaxing—or even eliminating—the strict dependence on MS1 feature detection, while preventing the generation of excessive false-positive pseudo-spectra. Another aspect worth exploring is the use of the correlation coefficient values computed during ion pairing. Although these values are currently used internally for precursor–fragment association, they are not yet propagated to the downstream DDA search engine. Leveraging such correlation metrics to refine scoring or confidence estimation could further enhance peptide identifications. In its current implementation, xTracer focuses exclusively on qualitative analysis and does not yet support peptide quantification. Future work will aim to construct spectral libraries from xTracer-derived identifications, enabling subsequent targeted quantification using library-based approaches. Moreover, the option in xTracer to compute correlations solely in the mobiligram dimension suggests potential applications to direct infusion–based HRIM workflows, where LC-based separation is minimal but ion mobility structure remains informative.

## 4 Conclusions

PAMAF represents a fundamentally new acquisition paradigm that achieves nearly complete ion utilization by fragmenting all mobility-separated precursors without quadrupole isolation. This approach maximizes ion sampling efficiency while preserving spectral specificity through the ultrahigh mobility resolution of SLIM, making PAMAF a highly efficient and analytically powerful platform for large-scale proteome profiling. Building upon this foundation, we developed xTracer, the first untargeted peptide identification algorithm designed specifically for PAMAF data. By integrating chromatogram- and mobiligram-based correlations, xTracer reconstructs pseudo-spectra directly from quadrupole-free, mobility-aligned fragmentation signals and enables confident peptide identification using well-established DDA search engines. This is impressive given the higher complexity of matching relatively low-resolution MS data at MS1 and MS2 data without simplification of fragment space by quadrupole selection. Applied across datasets with different sample loads and acquisition throughputs, xTracer consistently achieved robust and reproducible peptide identifications, outperforming single-domain correlation strategies. In summary, xTracer establishes a general computational framework for interpreting quadrupole-free, mobility-aligned fragmentation spectra, expanding the analytical potential of high-resolution ion mobility–based proteomics.

## Acknowledgements

This work was funded by NIGMS R35GM142502.

## Author contributions

JS, DD and JGM contributed to the conceptualization of the study. JS was responsible for data curation (together with LR and LD), formal analysis, methodology design (together with DD and JGM), software development (together with BK) and validation (together with LD). Funding acquisition and supervision were provided by DD and JGM. Resources were provided by DD. JS, DD and JGM wrote the original draft of the manuscript. All authors participated in the review and editing of the manuscript.

## Supplementary Information

**Supplementary Fig. 1.**
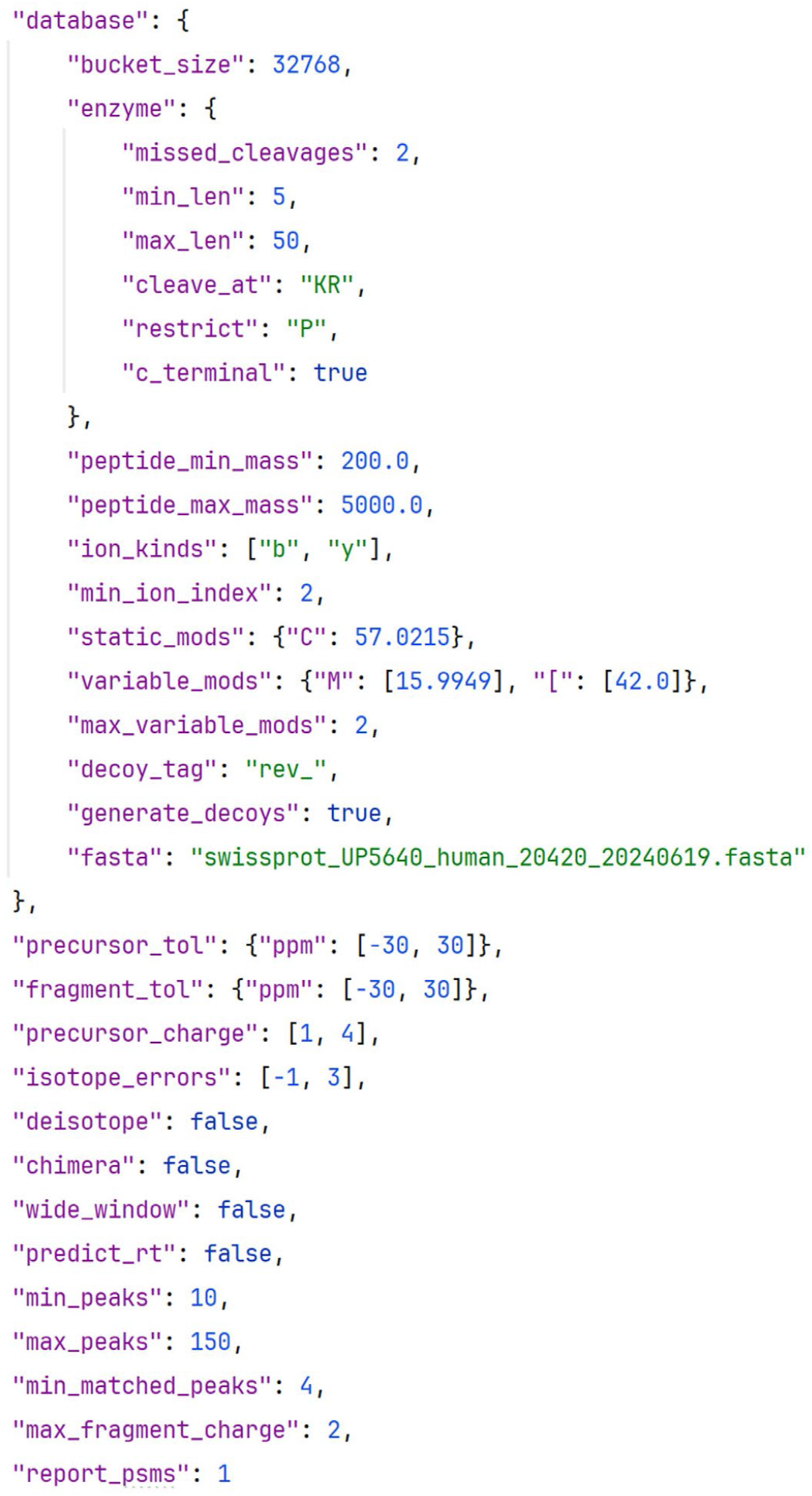
Configuration parameters of Sage. When the mass tolerance and minimum peak count parameters of xTracer are adjusted, the Sage parameters “precursor_tol”, “fragment_tol”, and “min_peaks” should change accordingly.

**Supplementary Fig. 2.**
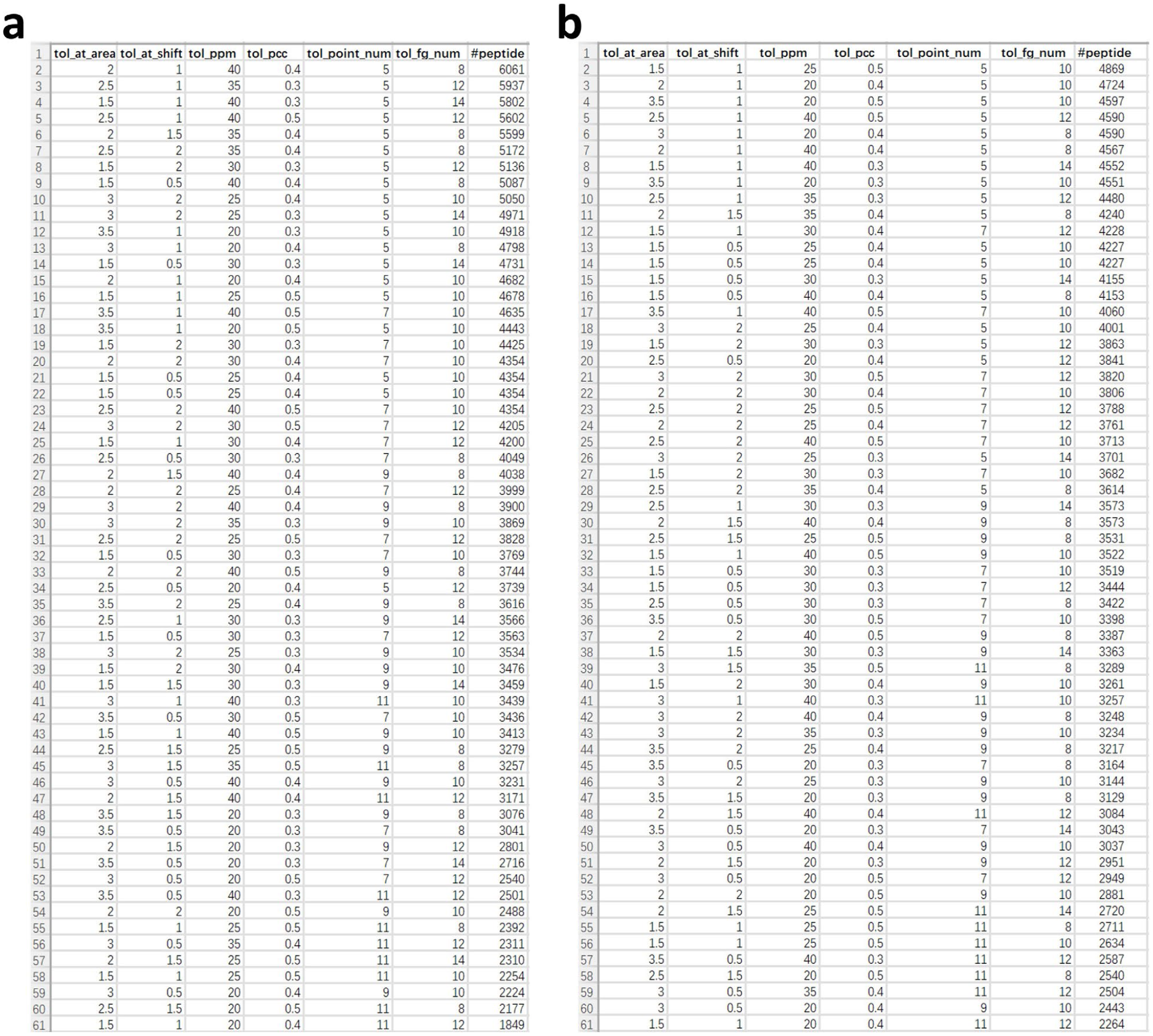
Number of peptide identifications under different xTracer parameter combinations. **a)** Data from the varying sample amount dataset, ‘2024-10-24 19.23.10-50ngHeLa_1iRT_CERamp-updated.mbi’. **b)** Data from the varying throughput dataset, 2024-10-31 01.37.30-100ng_HeLa_1irt_100S-updated.mbi’.

**Supplementary Fig. 3.**
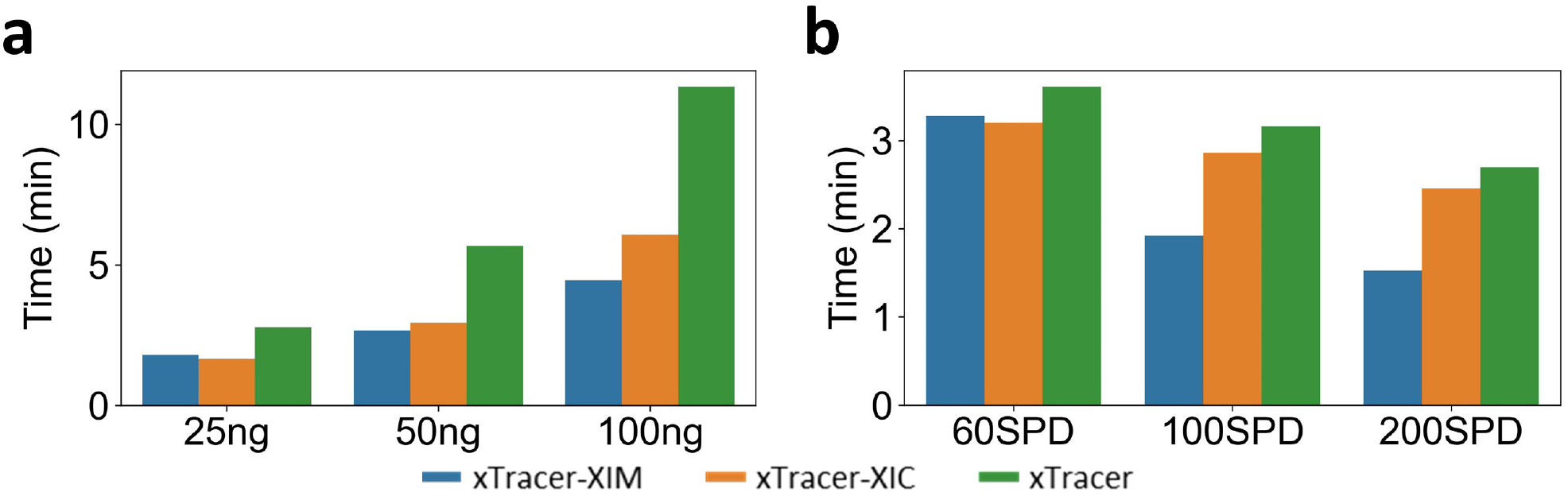
Evaluation of analysis time across. **a)** different sample loads and **b)** varying acquisition throughputs datasets.

